# Quantitative manipulation of the photoperiodic flowering by a small-molecule clock modulator

**DOI:** 10.1101/2025.11.06.686944

**Authors:** Akari E Maeda, Shogo Ito, Tokitaka Oyama, Ayato Sato, Norihito Nakamichi, Tomoaki Muranaka

## Abstract

Photoperiodism, a seasonal response mechanism that relies on day-length measurement by the circadian clock, is a major regulator of flowering time ^1^. The external coincidence model proposes that the alignment between the internal circadian phase and external light signals triggers photoperiodic responses ^2^. It has been challenging to establish robust models for the quantitative relationship between clock and flowering time, due to a lack of tools to modulate clock quantitatively. Here, using a circadian period-lengthening small molecule, we demonstrated the quantitative modulation of the critical day length for flowering in monocots.

## Main text

In *Arabidopsis thaliana* (Arabidopsis), expression of *FT*, encoding a florigen, is induced by CONSTANS (CO) which is expressed in the evening and stabilized by far-red and blue lights. Thus, a period mutant affects the phase of CO expression and flowering time under light-dark cycles ^3^. The relationship between the free-running period and phase for photoperiodic flowering was also found in a monocot ^4^. Thus, the manipulation of the circadian period possibly changes the photoperiodic flowering in a quantitative manner.

We previously reported that an inhibitor of Casein Kinase 1 (CK1), a highly conserved kinase involved in circadian period regulation in eukaryotes, lengthens the circadian period in Arabidopsis in a dose-dependent manner ^5,6^. *Lemna aequinoctialis* is a small floating monocot that exhibits rapid short-day flowering with high photoperiod sensitivity ^4^. Here, we modulate photoperiodic flowering in *L. aequinoctialis* using a CK1 inhibitor, B-AZ(3,4,-dibromo-7-azaindole).

To consider B-AZ targets in *L. aequinoctialis*, we used a deposited transcriptome (PRJDB12719) and identified six top candidate CK1 genes belonging to three phylogenetic clusters (Figure 1A). In Arabidopsis, 13 *CK1* homologs have been identified and are referred to as *CASEIN KINASE 1 LIKE* (*CKL*) genes. Among them, AtCKL4 has strong kinase activity of the clock proteins, and the activity was inhibited by B-AZ *in vitro* ^5^. Therefore, we selected the *L. aequinoctialis CK1* homolog belonging to the *AtCKL3*/*AtCKL4* cluster (hereafter *LaCKL*) as possible targets of B-AZ.

**Figure 1.**
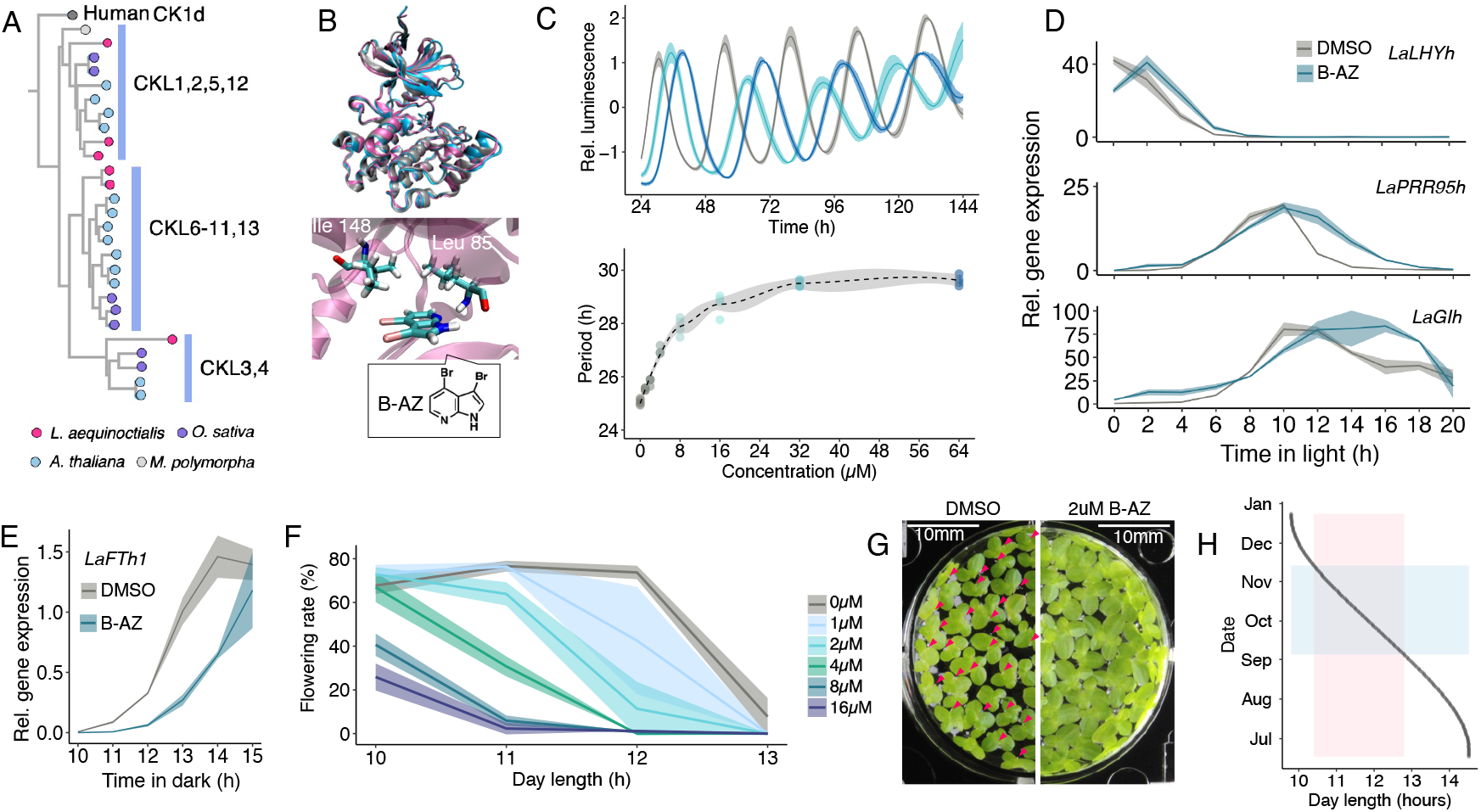
The CK1 inhibitor, B-AZ, quantitatively manipulates the circadian clock and photoperiodic flowering in *Lemna* plants. (A) Phylogenetic tree of CK1 homologs. Magenta-, blue-, purple-, white-, and gray-colored circles represent *L. aequinoctialis, A. thaliana, Oryza sativa, Marchantia polymorpha*, and *Homo sapiens*, respectively. (B) The superimposed image of predicted structures of LaCKL (1–292 aa, magenta), AtCKL (1–293 aa, cyan), and the crystal structure of human CK1δ (PDB: 5IH6) (top). Simulated binding structure of B-AZ in the ATP-binding pocket of LaCKL, with Leu85 and Ile148 residues spatially associated with B-AZ (bottom). (C) Luminescence rhythms of *AtCCA1:LUC* (top) and corresponding circadian period length (bottom) in *L. japonica* treated with various concentration of B-AZ (n = 4). The dashed line indicates the LOESS regression fit; the shaded area represents the 95% confidence interval. (D) Relative mRNA expression levels of clock genes *LaLHYh* (top), *LaPRR95h* (middle), and *LaGIh* (bottom) in *L. aequinoctialis* treated with DMSO (gray) or 8 µM B-AZ (blue) (mean ± SD, n = 3). (E) Relative mRNA expression of *LaFTh1* in *L. aequinoctialis* treated with DMSO (gray) or 8 µM B-AZ (blue) (mean ± SD, n = 3). (F) Photoperiodic responses of flowering in *L. aequinoctialis* treated with DMSO or B-AZ. Flowering rates after 9 days of photoperiodic treatment are plotted against day length (mean ± SD, n = 4). (G) *L. aequinoctialis* colonies treated with DMSO or 2µM B-AZ for a month. Flowers are pointed by magenta triangles. (H) Day length changes at latitude 35° N from summer solstice to winter solstice. The pink box indicates the range of critical day lengths that B-AZ can manipulate. The blue box indicates the range of days corresponding to the critical day length.

Protein structures of *AtCKL4* and *LaCKL* were predicted using ColabFold and showed high similarity to the crystal structure of human CK1d (Protein Data Bank ID; PDB: 5IH6), particularly within the kinase domain, indicating that LaCKL likely retains conserved kinase activity. Docking simulations confirmed that B-AZ binds to its ATP-binding pocket of LaCKL, suggesting that B-AZ inhibits the LaCKL (Figure 1B).

To assess the potential for chemical tuning of the circadian clock in *Lemna* plants, we first examined the effects of B-AZ on *L. japonica* 5512 carrying the circadian bioluminescence reporter *AtCCA1:LUC* ^7^. B-AZ lengthened the circadian period of *AtCCA1:LUC* reporter in a dose-dependent manner (Figure 1C). 1 µM, 4 µM, and 8 µM lengthened the period by 0.5 h, 1.9 h, and 2.8 h, respectively. The effect was saturated at 32 µM by 4.4 h.

Next, we analyzed the diel expression patterns of clock-related genes in *L. aequinoctialis* 5636. The homologs of three clock-related genes, *LATE ELONGATED HYPOCOTYL* (*LaLHYh), PSEUDO-RESPONSE REGULATOR 95* (*LaPRR95h*), and *GIGANTEA* (*LaGIh)*, were identified from the deposited transcriptome data and analyzed by reverse transcription quantitative PCR (RT-qPCR) (Figure S1 and Table S1) ^4^. Similar to Arabidopsis, the peak expression of *LaLHYh, LaPRR95h*, and *LaGIh* occurred at subjective dawn, day, and night, respectively (Figure 1D). Treatment with 8 µM B-AZ resulted in a phase delay of all three genes (Figure 1D). The B-AZ treatment delayed the peak phase of *LaLHYh* by at least 2 h. *LaPRR95* peak phase estimated by the fitting model suggested 1.1 h delay by B-AZ. Peak phase of *LaGIh* was delayed at least 4 h by B-AZ.

In *L. aequinoctialis*, one of the *FT* homologs, *LaFTh1*, was induced at night, and its induction timing was associated with the critical day length of photoperiodic flowering ^4^. In the control treatment (Mock, 0 µM B-AZ), the *LaFTh1* expression was induced by 11 h dark and saturated within 14 h dark (Figure 1E). In the 8µM B-AZ treatment, 13 h dark was required for *LaFTh1* induction. 15 h of dark was not sufficient for the full expression of *LaFTh1*. The phase delay effect on *LaFTh1* expression implied that B-AZ quantitatively changes photoperiodic flowering of L. *aequinoctialis*.

We treated *L. aequinoctialis* with different concentrations of B-AZ (0 ∼ 16 µM) at four daylength conditions (10, 11, 12, or 13 h light) (Figure 1F). In the control treatment (Mock, 0 µM B-AZ), flowering was fully induced under 10, 11, and 12 h light conditions, but poorly under 13 h light. In the 1 µM B-AZ treatment, although flowering was fully induced under 11 h light, flowering rate was reduced at the half level under 12 h light. In the 4 µM B-AZ treatment, flowering was not induced even under 12 h light conditions but fully induced under 10 h light conditions. B-AZ at 8 or 16 µM had stronger inhibitory effects on flowering. The estimated critical daylengths (defined as the daylength giving 50% of the maximum flowering rate) were 12.8, 12.0, 11.5, 10.9, 10.6, and 10.4 h for 0, 1, 2, 4, 8, and 16 µM B-AZ treatments, respectively. We also noticed that the treatment with 2 µM B-AZ under 12 h light maintained the plants in a vegetative state for over a month without detectable growth inhibition (Figure 1G). Collectively, B-AZ treatment shifted the critical daylength in a clear, dose-dependent manner. The observed 2.4 h shift corresponds to an approximately 70-day delay in flowering at a temperate latitude, 35° N (Figure 1H). Thus, only by manipulating the circadian clock period, the seasonal variation of flowering time be sufficiently achieved.

CKLs, the target of B-AZ, phosphorylate Arabidopsis core clock proteins, affecting their stability ^5,6^. Thus, post-translational regulations by CKLs may underpin determination of period length in duckweed. Given that *CO*/*Heading date 1* (*Hd1*) homologue was not found, the photoperiodic pathway in *L. aequinoctialis* may differ from that in Arabidopsis and in rice^4^. Despite these differences, the relationship between circadian period and flowering was evidenced both in short-day monocot and long-day dicot ^3,4^. Our study further highlights the quantitative manipulation of the circadian clock to regulate, modulate, and fine-tune output physiology crucial for agricultural applications ^8, 9, 10^.

## Acknowledgments

This work was supported by Japan Society for the Promotion of Science (JSPS) KAKENHI [Grant Numbers, JP23K14253 (T. M.), JP24H02304 (T. M.), JP24H02116 (T. M.), JP24H02121 (S. I.), and

JP25KJ0169 (A. E M.)], Japan Science and Technology Agency (JST) SATREPS (JPMJSA2004, T. M., S. I., and T. O.).

