## Supplemental information for "Quantitative manipulation of the photoperiodic flowering by a small-molecule clock modulator"

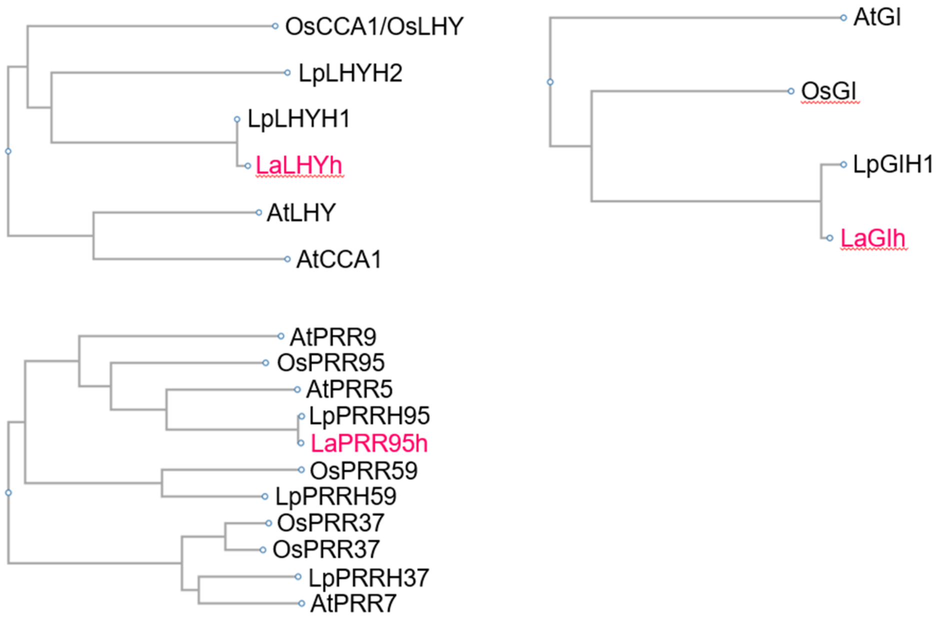


Figure S1.

Phylogenetic tree of *LHY, PRRs,* and *GI* homologs. OS: *Oryza sativa*, Lp: *Lemna paucicostata* (Synonym of *Lemna aequinoctialis*), At: *Arabidopsis thaliana*. The three homologs analyzed in this study are highlighted by red.

| qPCR primer name | Sequence | Source |
|---|---|---|
| LaFTh1_fwd | ACCCTACCCTTAGAGAATATCTGC | Muranaka et al., 2022, iScience |
| LaFTh1_rev | TAGGTGCTGGGCCTTCATAG | Muranaka et al., 2022, iScience |
| LaACT2_fwd | ACACAGTGCCCATCTATGAAGG | Muranaka et al., 2022, iScience |
| LaACT2_rev | AGTAGCCTCGTTCGGTTAGGATC | Muranaka et al., 2022, iScience |
| LaGIh_fwd | TGCCTGGATTACGGATATTCTC | This paper |
| LaGIh_rev | CTTCATCGCAGACACTCAGTA | This paper |
| LaPRR95h_fwd | ATGATTCTTGCAAGAGCATTCC | This paper |
| LaPRR95h_rev | GGTGTGTTCTCGGCTCCATT | This paper |
| LaLHYh_fwd | GGAAACCCTACACTATCACCAAAC | This paper |
| LaLHYh_rev | GGATGTCGATGTCGTGGACTTG | This paper |

Table S1. Primers for RT-qPCR

### Supplemental experimental procedures

#### Plant materials and growth conditions

Transgenic *L. japonica* 5512 and *L. aequinoctialis* 5636 (formerly called as Nd) were maintained in NT medium with 1% sucrose under 15-h light/9-h dark conditions (approximately 50 µmol $m^{-2}$ $s^{-1}$, white LED T5LT20W, Beamtec, Japan) in an incubator (HCLP-1280, NK system, Japan) at 25˚C. The NT medium is a modified version of NF medium, where $KNO_3$ and $KH_2PO_4$ were changed to 15 mM and 1.5 mM, respectively [S[1]].

#### Luminescence monitoring

Five to six colonies of *L. japonica* were transferred to a 40-mm polystyrene dish (Asnol Petri Dish, AS ONE, Japan) with 5 ml of growth medium (NT with 1% sucrose) containing 0.1 mM D-luciferin (Wako, Japan). The 1µl of dimethyl sulfoxide (DMSO) containing B-AZ was added to each diches. The luminescence from *L. Japonica* carrying *AtCCA1:LUC* was automatically monitored with a luminescence dish-monitoring system with photomultiplier tubes under the continuous light condition at 25˚C (H7360-01; Hamamatsu Photonics K.K., Japan) as previously (Muranaka et al., 2022). The period length was estimated by FFT-NLLS, as previously described [S[2]].

#### Phylogenic analysis of CK1 homologs

The transcript sequences of *L. aequinoctialis* were obtained by *de novo* assemble of deposited transcriptome (PRJDB12719) using TRINITY v2.15.2 [S[3]]. The amino acid sequences were then estimated using TransDecoder v5.7.1. The *CKL* homologs were searched by BLASTP with a query for the AtCKL4 amino acid sequence. The homologs of *A. thaliana*, *O. sativa*, and *M. polymorpha* were obtained by ORTHOSCOPE with a query for the AtCKL4 amino acid sequence [S[4]]. The amino acid sequence of human CK1d was obtained from Uniprot (Uniprot-ID: P48730). The phylogenetic tree was constructed using the online tool ClustalW (https://www.genome.jp/tools-bin/clustalw) with PhyML algorithm. As outgroup sequences, amino acid sequences of *A. thaliana* MUT9-LIKE KINASE1 (MLK1) and human CK1g (Uniprot-ID: Q9HCP0) were used. The AtMLK1 was detected as a homolog of AtCKL4 in ORTHOSCOPE analysis.

#### Comparison of protein structure

The protein structures of LaCKL and AtCKL4 were predicted by ColabFold v1.5.5 [S[5]]. To compare the structure between human CK1d (PDB ID: 5IH6,[S[6]]) and the predicted structures, whole protein structures were superimposed with UCSF Chimera v1.15 [S[7]] and visualized with VMD v0.0.1d1 [S[8]]. We utilized DynamicBind [S[9]] as a docking simulation between LaCKL and B-AZ.

**Gene expression analysis for *LaLHYh*, *LaPRR95h* and *LaGIh***

The *LaLHYh, LaPRR95h,* and *LaGIh* homologs were determined in the *de novo* assembled sequences by BLASTP with a query for the LpLHYH1 (GenBank: BAD97866.1), LpPRRH95 (GenBank: BAE72699.1), and LpGIH1 (GenBank: BAD97864.1) amino acid sequences, respectively [S10]. Amino acid sequences of the homologs of these genes were obtained by ORTHOSCOPE, and the phylogenetic tress of those were constructed by ClustalW as described for CK1 homologs.

*L. aequinoctialis* colonies were grown under 15-h light /9-h dark at 25˚C. Two colonies of *L. aequinoctialis* were transferred to a 40-mm polystyrene dish with 3 ml growth medium (NT with 1% sucrose) containing DMSO or 8μM B-AZ for 2 days. The dishes were then transferred into constant light. Colonies were collected in a tube with zirconia beads and flash-frozen in liquid nitrogen. Colonies were crushed with a Tissue Lyser II (QIAGEN), and the RNA was isolated by Illustra RNAspin (Cytiva). Reverse transcription followed by quantitative polymerase chain reaction for *LaLHYh*, *LaPRR95h*, *LaGIh* was performed with the normalization gene *LaACT2*. The qPCR primers for *LaLHYh*, *LaPRR95h*, *LaGIh* were designed from *de novo* assembled sequences and shown in Table S1.

**Gene expression analysis for *LaFTh1***

*L. aequinoctialis* colonies were grown under 15-h light /9-h dark at 25˚C. Three to four colonies of *L. aequinoctialis* were transferred to a 40-mm polystyrene dish filled with 3 ml of growth medium (NT with 1% sucrose) containing DMSO or 8μM B-AZ and cultured for 2 days under the same conditions. Then, they were released to constant darkness at the end of light, and sampled at 10, 11, 12, 13, 14, 15-h after release. The qPCR analysis was performed as mentioned above. The qPCR primers were shown in Table S1.

**Measurement of the critical day length in photoperiodic flowering**

A single colony of *L. aequinoctialis* was placed in each well of a 24-well plastic plate filled with 2 mL of growth medium (NT with 1% sucrose) containing DMSO or 1, 2, 4, 8, 16μM B-AZ. These plants were grown under the six conditions of day-length (9-h light/15-h dark, 10-h light/14-h dark, 11-h light/13-h dark, 12-h light/12-h dark, 13-h light/11-h dark) at 25˚C for 9 days. After the treatments, the plants were bleached with 70% ethanol. The number of flowering and non-flowering fronds were counted for each colony under a stereoscopic microscope. Flowering rate was calculated as the percentage of flowering fronds to the total number of fronds in four biological replicates.

**Growth analysis**

A single colony of *L. aequinoctialis* was placed in each well of a 6-well plastic plate filled with 10 mL of growth medium (NT with 1% sucrose) containing DMSO or 2μM B-AZ. These plants were grown under 12-h light/12-h dark conditions at 25°C. Every one week, one colony was transferred to new medium. After a month from treatment start, the photos were taken after 13 days culture from transfer.

**Day-length calculations**

Day lengths shown in Figure 1H were calculated using the online service of the National Astronomical Observatory of Japan (https://eco.mtk.nao.ac.jp/cgi-bin/koyomi/koyomix_en.cgi).
